# Topoisomerase Based Control of Cellular Transcription and Growth

**DOI:** 10.1101/2025.05.26.655193

**Authors:** Harris Clark, Cole Giusto, Raaghav Thirumaligai, Enoch Yeung

## Abstract

Global cellular transcription and growth are tightly coupled to the supercoiling state of a cell’s genomic DNA. Direct processes that modulate the supercoiling state of genomic DNA thus provide a novel mechanistic route for biological control of cellular dynamics. In this work, we develop a stochastic model of genome-wide transcriptional responses to local DNA supercoiling and show that, despite heterogeneity in the transcription of each individual gene, the overall system converges to a unique stationary distribution. From the stochastic system, we derive a corresponding nonlinear population model with both local and global supercoiling states, demonstrate that the proposed model exhibits a unique, globally stable equilibrium point, and identify key parameters through sensitivity analysis. We further extend the model by considering the temperature-sensitive gyrase mutant *E. coli* strain nalA43 and illustrate how shifting the gyrase–topoisomerase balance via temperature acts as an effective control knob on the population, from which we derive a control law. Numerical simulations confirm the effectiveness of the derived control law and provide insight into the dynamics of the system. These results help establish a theoretical foundation for understanding the crucial equilibrium dynamics of topoisomerase enzymes and their implications for cell function.

## I. Introduction

DNA supercoiling is a fundamental biological phenomenon that describes the relative increase or decrease in twist in the DNA double helix. Excessive twist can lead to rotation of the DNA double helix in 3D space, known as writhing, and facilitate compaction of DNA in concert with DNA binding proteins [1], [2], [3]. Proper regulation of supercoiling is crucial for transcription, as DNA topology directly affects the accessibility and activity of the transcriptional machinery [1], [2], [4], [5], [6].

Around 70-80% of the supercoiling in the cell is the direct result of transcription (The rest as a result of DNA replication) [7], controlling the transcription induced supercoiling will be the focus of our paper. During the transcription of a single gene, torsional stress is induced in the DNA when RNA polymerase unwinds the DNA helix, opening up a transcription bubble, and twisting it into supercoiled domains [1], [5], [6], [8]. The complexity of DNA supercoiling arises when structures such as plectonemes, form when the double helix twists in three-dimensional space to relieve the additional torsional stress [9], [3]. These properties contribute to transcriptional bursting, in which genes cycle between active and inactive states due to accumulated torsional stress [4], [5]. Such dynamic regulation makes supercoiling an essential cellular control mechanism, impacting cellular behavior, genetic stability, and adaptation [10], [5], [6].

In synthetic biology, manipulating DNA supercoiling offers a promising strategy for controlling genetic circuits and engineering gene expression [10], [11]. Unlike traditional transcription factor or CRISPR based control mechanisms, supercoiling acts as a near instantaneous feedback mechanism, not limited by the slower rates of protein translation and folding. Precise modulation of DNA supercoiling could enable the creation of genetic systems with more predictable and tunable behaviors [10], [11]. In this work, we explore the biophysical mechanisms underlying DNA supercoiling and investigate transcriptional control achieved by creating an imbalance between DNA gyrase and topoisomerase activity using the thermally sensitive *E. coli* mutant nalA43 [12]. Understanding these mechanisms facilitates the design of more sophisticated and robust engineered biological systems.

## II. topoisomerase Based Population Dynamics

In order to develop a comprehensive model of cell population dynamics that accounts for the effects of topoisomerase enzymes, we model our system from the bottom-up. First, we derive the individual dynamics of transcription bubbles, DNA supercoiling, topoisomerases, and mRNA. We then integrate these dynamics to create a novel class of cell population models

As a starting point, we focus on the dynamics of transcription bubble formation across the cell. We consider a simple chemical reaction in which we assume that the catalytic rate of formation, *k*_*cat*_, is dependent on the global average supercoiling of the DNA, *σ*, and that the bubbles decay at a first order rate.

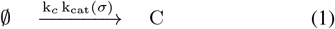

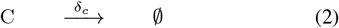

Where *C* represents a complex of the transcription bubble and its associated polymerase enzymes, *k*_*c*_ is the rate of formation, *δ*_*c*_ is the rate of collapse, and *k*_*cat*_(*σ*) models the effects of the supercoiling-dependent rate of initiation.

We assume that the rate of formation *k*_*cat*_(*σ*) has the functional form of a radial basis function [10], [11], [13], [14], which matches the empirical observations in [15] and [16]. We fit this function as:

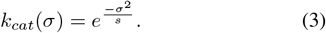

Throughout this paper we assume that the coordinates of the global supercoiling density are centered around the homeostatic setpoint *σ*^∗^ = 0.095, equivalent with the 10.5 basepairs per turn in standard B-form DNA [2], [11]. When we write the state variable *σ* in this paper, it will correspond to the absolute supercoiling density.

From this network we can develop an ODE that describes our bubble dynamics from the law of mass action [17]:

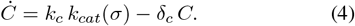

There are thousands of genes in a cell’s genome, of which hundreds are being transcribed at any given moment. Each gene has unique length scales and properties that contribute to its distinct local supercoiling state. When a specific gene is transcribed, a transcription bubble forms, partially unwinding the DNA around the coding sequence. Due to DNA’s helical structure, this unwinding induces torsional stress, creating two distinct supercoiling domains [1], [8]. Ahead of the bubble (in the 5’ → 3’ direction on the coding strand), DNA becomes positively supercoiled, generating additional right-handed twists that hinder further unwinding. Behind the bubble (in the 3’ → 5’ direction), DNA becomes negatively supercoiled, forming left-handed twists [1], [8]. This localized supercoiling influences transcription efficiency and requires regulation by topoisomerase enzymes to maintain genomic stability [4]. In our model, we consider two topoisomerase enzymes: Type I topoisomerase (henceforth referred to as just ‘topoisomerase’), which relaxes negative supercoils, and Type II topoisomerase (referred to as ‘gyrase’), which relaxes positive supercoils.

**Fig. 1.**
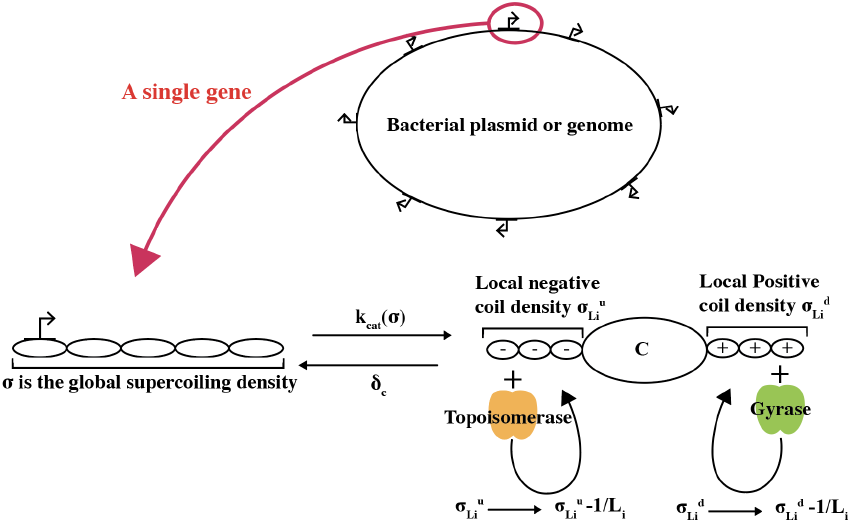
A schematic illustrating the region of interest. We model how the local supercoiling dynamics effect the global supercoiling level.

Each transcription bubble is characterized by two continuous variables: the positive local supercoiling density (upstream of the transcription bubble), 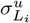,and the negative local supercoiling density (downstream of the transcription bubble), 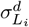.These densities represent the number of coils per unit length over a local domain of length *L*_*i*_. Every time a bubble is opened it is injected with fixed initial supercoiling densities determined by the bubble’s size. Because transcription bubbles vary in size, their initial supercoiling densities and local length scales are heterogeneous, necessitating explicit modeling of bubble size variability to later understand the genome wide effects of local transcription.

First, whenever a bubble is opened, its local supercoiling density is fixed to some initial coil density, which is determined by dividing the discrete amount of injected coils by the local length.

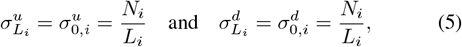

where *N*_*i*_ is the number of coils introduced when the transcription bubble for gene *i* is opened, and *L*_*i*_ is the length of the local region.

Over a bubble’s lifetime, the local densities decrease continuously due to enzyme-mediated degradation: gyrase removes positive supercoils at a rate proportional to *γ*_1_ *G*, and topoisomerase removes negative supercoils at a rate proportional to *τ*_1_ *T*, where *G* and *T* are state variables corrosponding to gyrase and topoisomerase, while *γ*_1_ and *τ*_1_ are the respective local-scale activity constants (both scaled by 1*/L*_*i*_ to convert from discrete coil numbers to a continuous coil density). Finally, when the bubble collapses at a rate *δ*_*c*_, its local state is completely removed. In the full system, bubble closure occurs at a rate *δ*_*c*_ *C*, where *C* is the number of bubbles.

To model this process while preserving gene heterogeneity, we use a chemical master equation followed by moment-generating functions. We begin by deriving a standard chemical master equation of our system.

Let 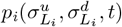 be the probability density that a bubble in gene *i* has local densities 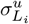 and 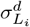 at time *t*. The CME for a single gene then consists of three components.

First, the injection term, which represents bubble opening at a rate defined in (4), is modeled by delta functions, which pin the concentrations when bubbles are formed at the fixed densities 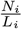:

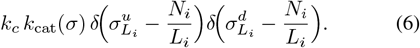

Second, the drift terms capture the continuous degradation of the densities. A generic drift term *ν* is introduced, as probability must be conserved, any change in the density is given by the negative divergence of that flux, such that:

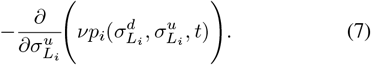

We model the relaxation of positive supercoils as a chemical reaction in which gyrase binds to a single local positive supercoil *N*_*i*_, and annihilates it. By converting this first order reaction to continuous form by dividing *N*_*i*_ by the local length *L*_*i*_ we can write this reaction as:

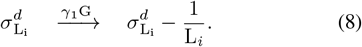

We can then derive the deterministic form of this contributing drift term from the law of mass action [17]. This captures the relaxation of positive coils as:

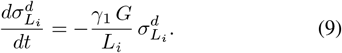

Thus, from the deterministic drift velocity, we can write it in probabilistic form as:

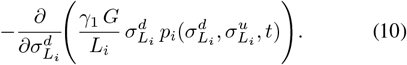

By analogy, the loss in negative density is represented as:

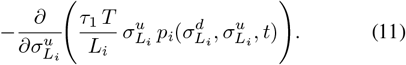

Finally, the closure term accounts for bubble collapse at a rate given by (4) and appears as a sink:

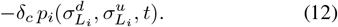

Thus, the complete CME for gene *i* is given by

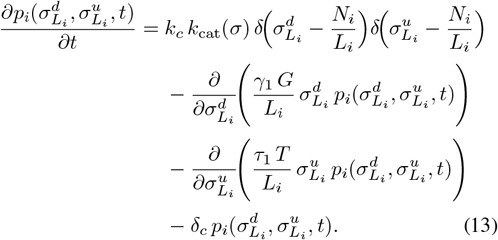

To derive the mean dynamics for gene *i*, we define the joint moment generating function

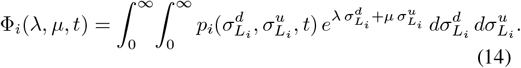

To extract the mean positive supercoiling density for gene *i*, we differentiate Φ_*i*_ with respect to *λ* and set *λ* = *µ* = 0 [17]:

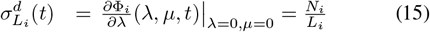

Thus, the injection term contributes 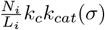.

By performing analogous computations for the negative injection term, drift terms, and sink terms [17] we derive the mean-field ODEs that describe the local positive and negative supercoiling dynamics for a single gene *i* as:

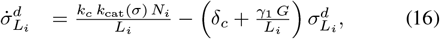

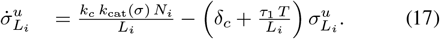

We introduce an additional term that defines the net local supercoiling density for gene *i* as 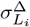.When a transcription bubble collapses, the positive and negative supercoiling domains interact, annihilating each other. However, if there was an initial imbalance between these domains prior to collapse (due to unequal gyrase and topoisomerase activity), a net contribution of supercoils remains and is transferred to the genome-wide supercoiling state. We model this net local supercoiling density as:

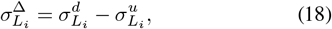

and by subtracting the above ODEs we have

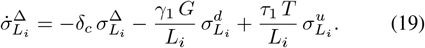

Now suppose that the genome contains *n*_*g*_ genes, each with its own local domain length *L*_*i*_ and injection parameter *N*_*i*_ (for *i* = 1, …, *n*_*g*_). For each gene *i* the dynamics are given by the CME in Eq. (13) and the corresponding mean-field ODEs. Assuming that the genes act independently and are completely isolated from neighboring supercoiling effects, the aggregate behavior is obtained via the product of the individual moment generating functions.

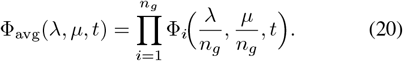

Differentiating Φ_avg_(*λ, µ, t*) with respect to *λ* and *µ* at *λ* = *µ* = 0 recovers the arithmetic means, corresponding to the average of a single gene:

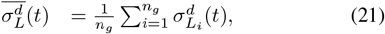

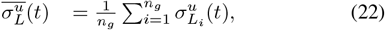

from which we can derive the mean local net density change:

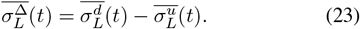

and then the mean total net change 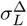:

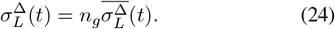

Finally we can derive the evolution of the global super-coiling due to net local by transferring the net local density to the global state. If the local densities are defined over domains with an average length ⟨*L*⟩ and the genome has total length *L*_*g*_, then the global supercoiling evolves as

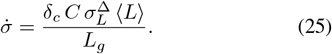

In addition, gyrase and topoisomerase also regulate the global supercoiling density. However, modeling these dynamics through a reaction network is challenging unless negative and positive global supercoiling are tracked separately. Since typical steady state values of *σ* in nature are only around − 0.06 [3], we introduce two exponential terms (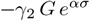 and 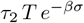 where *α* and *β* are binding sensitivity parameters) that are good approximations of the binding kinetics between *σ* and *G* and *T* which would be found from a typical mass action approximation. This approach keeps our system analytically clean and continuous, avoiding piecewise solutions. We are able to define 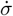 as:

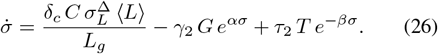

Having derived how the global supercoiling state *σ* and the levels of gyrase (*G*) and topoisomerase (*T*) evolve, we now connect these gene-level dynamics to cell population growth. In many microbes, the rate at which new cells are produced depends strongly on their capacity to transcribe and translate essential genes. Since the transcription rate itself is regulated by supercoiling, supercoiling is therefore able to modulate overall growth.

Thus, we consider a standard model for supercoiling-influenced transcription of some arbitrary gene *M* :

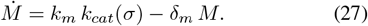

We apply the same form to both gyrase and topoisomerase, ignoring the time delay from translation for simplicity:

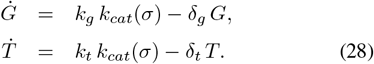

We can now consider a standard model for logistic population growth of some population *P* :

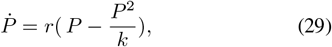

where *r* is a constant corresponding to the growth rate and *k* is the carrying capacity. Given that a cell’s growth ultimately depends on gene transcription, for our purposes *r* is not constant, but rather a function of *σ* that relates to the rate of transcription—*k*_*cat*_. We then replace *r* with *r k*_*cat*_(*σ*) to account for this dependency. As such we write our model for supercoiling-dependent population growth as

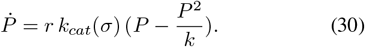

Thus, the population increases monotonically when *P* is less than its carrying capacity *k*, but the supercoil dependency means that at high levels of supercoiling in the cell, growth rate is arrested.

We have derived our complete system of 7 coupled ODEs:

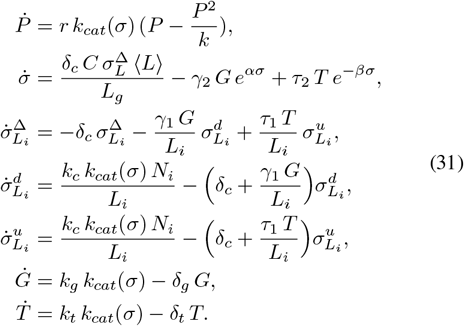

### Proposition 1

The system (31) has a unique, nontrivial equilibrium point

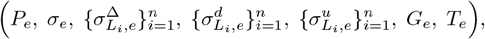

given by

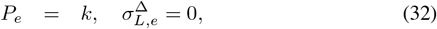

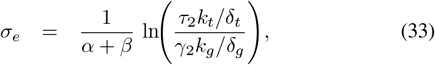

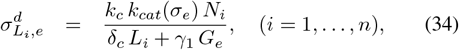

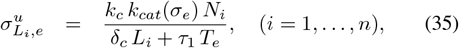

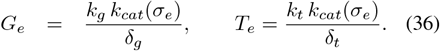

*Proof:* We determine the equilibrium of the system by setting

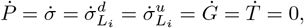

for each gene *i* = 1, …, *n*_*g*_.

### Local Dynamics

For the positive local supercoiling density in gene *i*, the mean-field ODE is

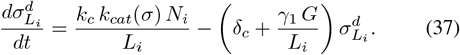

At equilibrium (with *σ* = *σ*_*e*_ and *G* = *G*_*e*_), we obtain

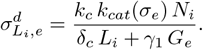

Similarly, for the negative local density we have

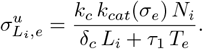

The equilibrium of the net local density is given by

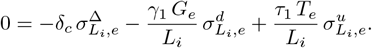

A sufficient condition is to choose 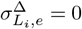,so that 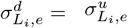. This, in turn, requires that

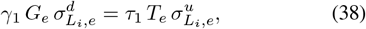

This physically means that there is no net local supercoiling when there is balanced Topoisomerase and gyrase activity. This is mathematically satisfied when the expressions in (34) and (35) are equated.

### Enzyme Dynamics

Setting 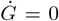 and 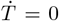,yields the following equilibria

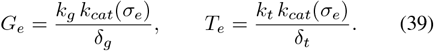

### Global Supercoiling Dynamics

The global supercoiling evolves according to

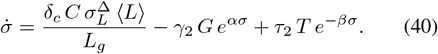

At equilibrium, since 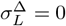 then from (40) we have

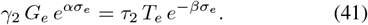

Taking the natural logarithm of both sides of (41) leads to:

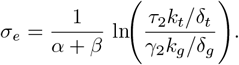

### Population Dynamics

The logistic population growth equation with supercoiling-modulated growth is

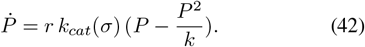

Setting 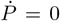 and discarding the trivial solution *P*_*e*_ = 0 yields

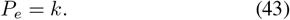

Thus, each equilibrium variable is uniquely determined, and the system possesses a unique, nontrivial equilibrium. ▪

## III. Sensitivity Analysis

By quantifying how sensitive the equilibrium points are to variations in specific parameters (such as gene length scales or enzyme production rates), we can determine which factors have the strongest influence on supercoiling homeostasis, enzyme balance, and overall population behavior. This, in turn, highlights the most important parameters for controlling the system’s dynamics. In this section we will investigate how the local domain lengths *L*_*i*_ and the rate constants *k*_*g*_ and *k*_*t*_ effect the systems equilibrium. *L*_*i*_ defines the density of the local positive and negative supercoiling domains, central to the heterogeneity of the model. In addition we analyze the variation of *k*_*g*_ and *k*_*t*_, the production constants of gyrase and topoisomerase, as these enzymes establish the careful equilibrium of supercoiling and underpin the control strategy presented in Section IV.

Using the explicit equations for the equilibrium values, sensitivity can be computed symbolically by taking derivatives of these values with respect to each parameter of interest [18]. This analysis provides insight into the model’s robustness to parametric uncertainty and identifies the parameters that most significantly influence equilibrium behavior.

For the following calculations, the equilibrium states are defined as 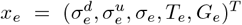.The sensitivity is calculated for the critical parameters *L*_*i*_, *k*_*g*_, and *k*_*t*_.

To simplify the local equations, we will use *C*_*g*_ to represent 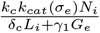 and

For each gene, the dependence of the equilibrium on *L*_*i*_ is shown below:

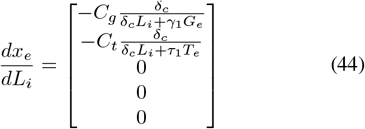

The zeros corresponding to the *σ*_*e*_, *T*_*e*_, and *G*_*e*_ rows show that changing the length of the local domain does not affect the values of *σ*_*e*_, *G*_*e*_, or *T*_*e*_. Increasing *L*_*i*_ only changes the equilibrium values for the local densities around the gene.

Below, the sensitivity of *k*_*g*_ and *k*_*t*_ is shown:

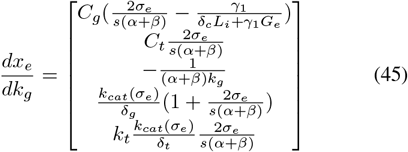

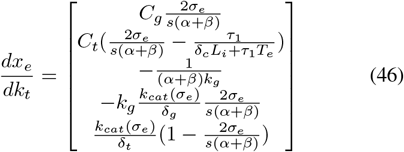

As expected, increasing *k*_*g*_ and *k*_*t*_ directly raises the equilibrium concentrations of gyrase (*G*_*e*_) and topoisomerase (*T*_*e*_), respectively. Consequently, an increase in *k*_*g*_ lowers the equilibrium global supercoiling density (*σ*_*e*_), while an increase in *k*_*t*_ elevates it. These effects are mediated by changes in the supercoiling-dependent transcription rate *k*_*cat*_(*σ*_*e*_), establishing a regulatory feedback loop that prevents excessive supercoiling. Specifically, at high supercoiling levels, expression of both gyrase and topoisomerase decreases, thereby reducing the net local supercoiling density 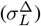 and stabilizing *σ*_*e*_.

## IV. Thermally Sensitive Mutants

*E. coli* strains with a nalA43 mutation display thermally sensitive gyrase activity [12]. A nalA43 mutation is a silencing mutation on the 43rd amino acid on the nalA gene, which encodes for the gyraseA subunit. This silencing mutation changes the structure of the gyraseA subunit such that it becomes sensitive thermal fluctuations, when grown at 42°C it was shown in the literature that the strain displays a 50-fold deficiency in this subunit [12]. This deficiency leads to a collapse of the colony population [12], as the cell loses its ability to regulate its supercoiling equilibrium [19]. In this section, we model this temperature-sensitive *E. coli* strain and show that increases in temperature shift the equilibrium population by altering the equilibrium global supercoiling density.

In order to capture the “turning off” of the activity of gyrase production between in 37°C and 42°C in the mutant strain nalA43 [12], we use a modified version of *k*_*g*_ that has the form of a logistic equation, such that there is a sharp down regulation of gyrase production between 37°C and 42°C:

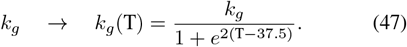

Note that the constants 2 and 37.5 were chosen as fit parameters, such that at 37°C *k*_*g*_(T) ≈ *k*_*g*_ and at 42°C the gyrase production is essentially negligible.

### Proposition 2

Assume that *k*_*g*_ in system (31) is replaced by its logistic form.

Then the unique, nontrivial equilibrium 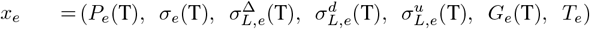 of the system retains the same structural form as in the constant-rate case.

*Proof:* Since every logistic-type rate constant *k*_*x*_(T) is strictly positive, the algebraic equilibrium conditions derived from

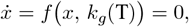

remain valid when *k*_*g*_ is replaced by *k*_*g*_(T). Thus, by the same algebra as before, the equilibrium retains the same form as (32) – (36), with *k*_*g*_ simply replaced by its temperature-dependent form *k*_*g*_(*T*). ▪

### Theorem 1

*(Temperature Sensitivity of Growth Rate):*

Consider the cell growth model

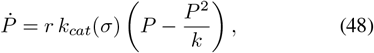

with the constraint that in the temperature sensitive system, the total gyrase activity is less than or equal to the total topoisomerase activity (*k*_*g*_ ≤*k*_*t*_, *γ*_1_ ≤*τ*_1_, *γ*_2_ ≤*τ*_2_) such that *σ* is always positive. Then, in the exponential growth phase (*P < k*) the temperature sensitivity

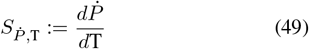

is strictly negative, while in the stationary phase (*P* = *k*) we have

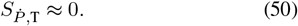

That is to say, increasing temperature can only slow or pause cell growth.

*Proof:* We begin with the growth rate (48). Although *k*_*g*_ is governed by a dynamic equation, its dependence on the effective gyrase production rate *k*_*g*_ remains analogous to the equilibrium case. Thus, *σ* satisfies

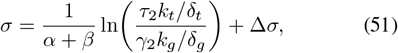

where Δ*σ* represents the non-equilibrium deviation. In order to account for the small deviations that occur as the temperature changes, we linearize *σ* about the equilibrium, so that the sensitivity of *σ* to *k*_*g*_ is given by

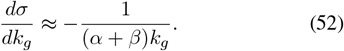

Then, differentiating *k*_*cat*_(*σ*) with respect to *k*_*g*_ via the chain rule gives

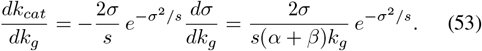

Differentiating 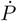 with respect to *k*_*g*_ yields

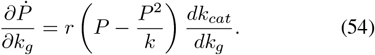

In the growth phase (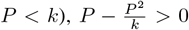,and from (53) we have 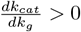,so that

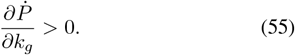

Next, the temperature dependence of *k*_*g*_ is modeled by (47), with derivative:

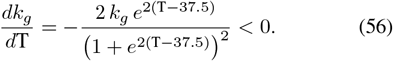

By the chain rule, the temperature sensitivity of the growth rate is:

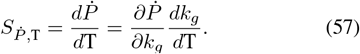

Since (55) implies 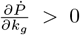 and (56) gives 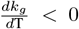, it follows from (57) tha

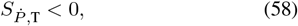

in the exponential growth phase.

In the stationary phase (*P* = *k*), so that

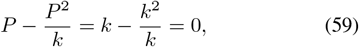

which immediately yields:

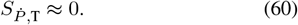

Thus, an increase in temperature can only slow or pause cell growth. ▪

With these results, we have the necessary mathematical conditions to develop a control law that relates cell population to culture temperature. This could have practical applications in biomanufacturing, providing a method to dynamically control the optical density of a bioreactor by manipulating its temperature in a reversible, non-invasive, and tunable manner.

### Theorem 2

Given the bounded reference signal *P*_*r*_(*t*), the full system is asymptotically stable under the dynamic control law:

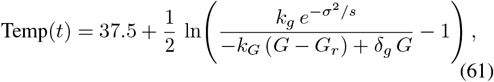

where Temp(t) denotes the controlled temperature of the ambient environment of a given reactor or culture of E. coli NalA43 cells.

*Proof:* Following the method of Sontag’s formula [20], to ensure that *P* tracks a desired reference *P*_*r*_(*t*), define the tracking error *e*_*P*_ (*t*) = *P*_*r*_(*t*) − *P* (*t*), and its derivative 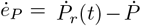. We enforce exponential decay by imposing 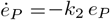. Since 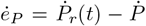,we can equivalently write this as 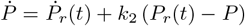.Equating this with the actual population dynamics we derive:

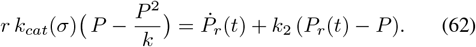

Solving for *k*_*cat*_(*σ*) yields:

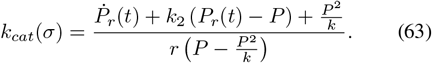

Define the effective catalytic rate as:

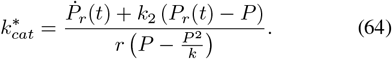

Since 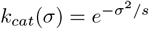,we can invert the equation for our desired global supercoiling reference *σ*_*r*_, with the convention that if 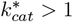,we set *σ*_*r*_ to 0, which is the maximum with *k*_*cat*_ = 1.

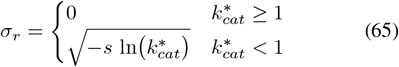

The derivative of *σ*_*r*_ is large. If the *σ* dynamics are assumed to be slow it can be left as 0. If desired, the derivative may be calculated, resulting in:

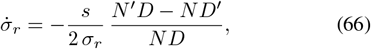

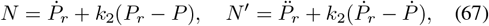

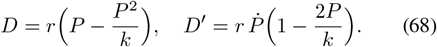

To ensure that the global supercoiling *σ* tracks *σ*_*r*_, we define additional error tracking terms *e*_*σ*_ = *σ* − *σ*_*r*_, and impose exponential decay with 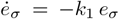.Thus, 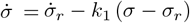. As above, we equate this with the actual supercoiling dynamics

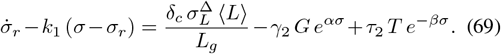

We can then solve for *G*, and define *G*_*r*_ as the desired gyrase level:

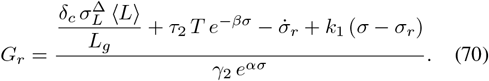

The gyrase production rate is modified to its logistic temperature dependence such that the effective production term becomes:

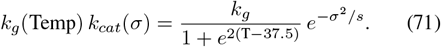

To force *G* to track *G*_*r*_, we impose

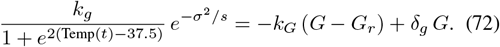

A direct algebraic manipulation yields

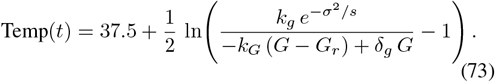

▪

## V. Simulation Results

In this section, we present numerical simulations of a diagnostic simulation that highlights the internal dynamics during a temperature-induced shift in gyrase-topoisomerase activity and our controlled thermally sensitive model (described in Section IV). The simulations were performed using Python 3.7 with the `scipy.odeint` solver on an M3 chip. The simulation code is available upon request.

### A. Mutant Gyrase-Topoisomerase Compensation

Gyrase and topoisomerase work together to maintain balance in the genomes supercoiling state. In work by [19], it was noted that, although gyrase is an essential enzyme, when gyrase was experimentally down regulated, some cell populations were still viable. Upon close inspection the only colonies remaining had a compensatory mutation in their topoisomerase mutation. It was hypothesized that this is because down regulating gyrase breaks the tight balance of supercoiling in the cell, causing supercoiling to shift to an unviable level which can only be restored with a compensatory mutation [19].

We show that our model corroborates these findings, we simulate two strains of nalA43 [12], one of which is defined as a “mutant” strain such that at t=400 minutes when the heat shock is applied, *k*_*t*_ is dropped, lowering the rate of topoisomerase production, simulating the compensatory mutation. We see that only the mutant strain is able to maintain viability after the heat shock. The simulations show that only the mutant strain maintains viability after the heat shock, with a smaller difference in local supercoiling densities leading to a lower final global supercoiling density compared to the wild type.

### B. Dynamic Population Control

We consider the controlled system given by (31) with the thermally sensitive *k*_*g*_ given in (47) with the control law from (73). As a theoretical challenge we attempted to simulate two equilibrium populations, by using a logistic reference population function defined as: 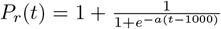 with its analytical first derivative: 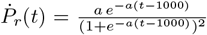 and second derivitave: 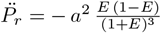.where *E* = exp(−*a*(*t* − 1000), and *a* = 0.05. The dynamics of the simulation are plotted in Figure 3. Consistent with Theorem 2, we see that the population is able to track our reference signal over a period of 2000 minutes. There are small perturbations, particularly with *σ* as the controller attempts to correct the population increase. Notably, given the model we chose for small timescales, there is no cell “death”, and as such the controlled rapidly adjusts to meet the reference signal, causing small oscillations.

**Fig. 2.**
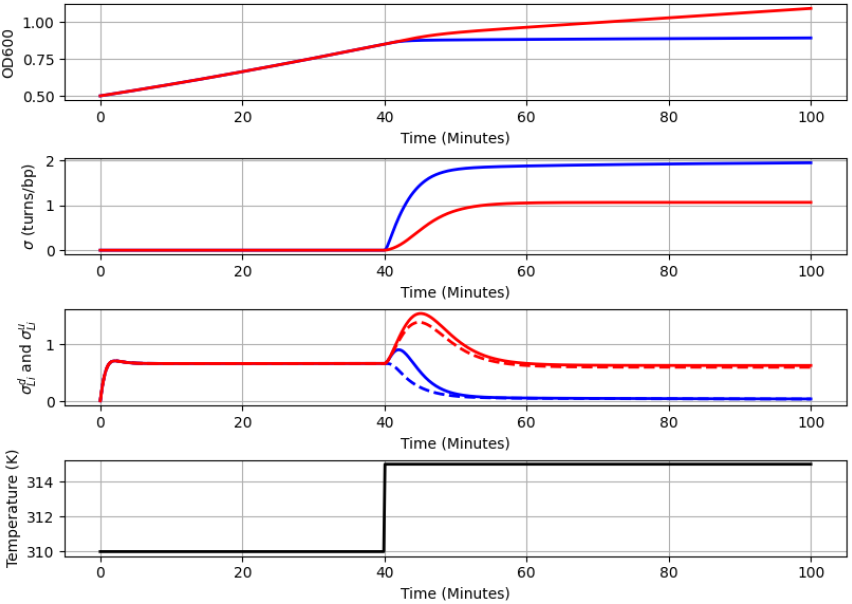
Population simulation of two “strains” of nalA43, one mutant (red) and one non-mutant (blue). There is a temperature shock at t=400 min which switches the temperature. The first subplot shows the OD600 over time. The second shows the global supercoiling density. The Third shows the local supercoiling densities, in which the solid line represents 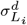,and the dotted line 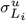.The last shows the temperature profile.

**Fig. 3.**
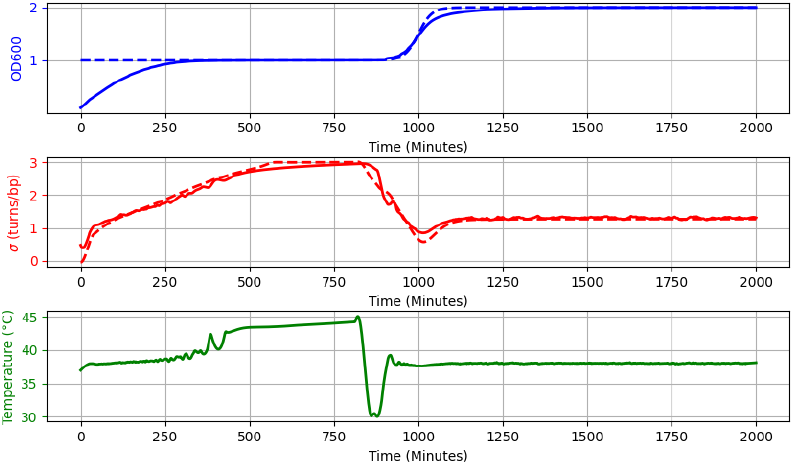
Dynamic tracking of a logistic reference population by modulating temperature. The system displays the ability to regulate the equilibrium population through temperature changes. The blue line represents the population, the red line represents the global supercoiling density, and the green line represents the temperature. The dotted line on the population graph represents the reference population, and the dotted line on the supercoiling graph is the desired supercoiling.

Implementing this controller in practice would require real-time measurements of *P, G, T*, and *σ*. Population *P* can be monitored via optical density (OD) measurements, with its derivatives computed computationally. Levels of *G* and *T* can be obtained using fluorescent markers and subsequent analysis, while high-resolution proxy measurements for *σ*(*t*) could be achieved through DNA visualization techniques which we anticipate having access to.

## VI. Acknowledgments

This work was funded in part by an NSF CAREER Award 2240176, the Army Young Investigator Program Award W911NF2010165 and the Institute of Collaborative Biotechnologies/Army Research Office grants ICBT-12YE01, W911NF19D0001, W911NF22F0005, W911NF190026, and W911NF2320006. This work was also supported in part by a subcontract awarded by the Pacific Northwest National Laboratory for the Secure Biosystems Design Science Focus Area “Persistence Control of Engineered Functions in Complex Soil Microbiomes” sponsored by the U.S. Department of Energy Office of Biological and Environmental Research. In addition we would like to thank Annie Nguyen, Yanran Wang, Lili Yang, Yizhuo Yin, Paige Nickerson, and Jai Mehra for their role in developing our understanding of supercoiling and how to model it.

